# ProtAug: An Empirical Investigation of pLM-Guided Data Augmentation for Protein Sequence Prediction Tasks

**DOI:** 10.64898/2026.07.10.737545

**Authors:** Zhuoyang Chen, Ruoqi Wang, Qiong Luo

**Affiliations:** Data Science and Analytics Thrust, The Hong Kong University of Science and Technology (Guangzhou), Guangzhou, China; Department of Computer Science and Engineering, The Hong Kong University of Science and Technology, Hong Kong, China

## Abstract

Protein language models (pLMs) offer great potential for protein sequence analysis, yet the scarcity of labeled data often limits their effectiveness in fine-tuning. Data augmentation is a promising remedy, but systematic evaluation of augmentation strategies for protein sequences remains limited, and the conditions under which augmentation confers downstream benefits are not well understood. In this paper, we systematically investigate pLM-guided substitution-based augmentation across seven protein prediction tasks. We propose ProtAug, a framework that leverages encoder-based (ESM-2) and autoregressive (ProtGPT2) pLMs to generate augmented sequences with user-controlled variation levels. Our investigation focuses on four questions: (Q1) whether pLM-synthesized sequences preserve more original signals than simpler methods, (Q2) to what extent augmentation improves prediction performance, (Q3) how variation levels affect downstream accuracy across tasks and models, and (Q4) whether biological plausibility is a necessary condition for achieving improvement. Our experimental results show that: (1) ProtAug Esm generally preserves motifs and structural similarity better than simple substitution, often comparable to homology retrieval; (2) augmentation yields consistent but task-dependent improvements, with ProtAug Esm achieving the best or second-best performance in 5 out of 7 tasks at 10% variation; (3) low-to-moderate variation levels (2–30%) perform best overall, although high-variation augmentation can benefit certain structure-related tasks; (4) the necessity of biological plausibility is task- and variation-dependent—while semantic preservation correlates with performance at low-to-moderate variation levels, improved generalization at high variation levels suggests that regularization effects, rather than label preservation, can also drive performance gains.

## 1 Introduction

Proteins are composed of sequences of amino acids that fold into three-dimensional structures. They play critical roles in various biological processes, such as structural support, molecular transport and catalysis. Similar to natural languages, where arrangements of words in sentences and paragraphs convey meanings, the sequences and structures in proteins contribute to protein functions and properties, or protein semantics.

As large language models have been widely applied in text-related tasks [Zhou *et al*., 2024], protein language models (pLMs) have been trained to generate semantic representations for protein sequences [Elnaggar *et al*., 2021; Rives *et al*., 2021]. These models, fine-tuned with labeled data, have demonstrated promising performance in various biological tasks. However, labeled protein data are scarce and expensive to acquire [Luck *et al*., 2020], especially those related to function, mutation, immune response, and interactions, or in rare species. Without sufficient and representative labeled data, pLMs may struggle to adapt to specific tasks. In this paper, we study how to effectively augment protein sequences to improve pLM performance in tasks.

Data augmentation has been used in both image processing [Shorten and Khoshgoftaar, 2019] and text processing [Wei and Zou, 2019]. In contrast, protein sequence augmentation remains largely unexplored. Existing augmentation strategies that perturb protein sequences, such as random substitution or shuffling, are often context-agnostic and may disrupt the structures and functions essential for effective augmentation (Figure 1). Recently, Sun et al. [Sun *et al*., 2024] conducted the first comprehensive benchmark of protein augmentation methods and proposed semantic-level strategies based on Integrated Gradients and back translation.

**Figure 1.**
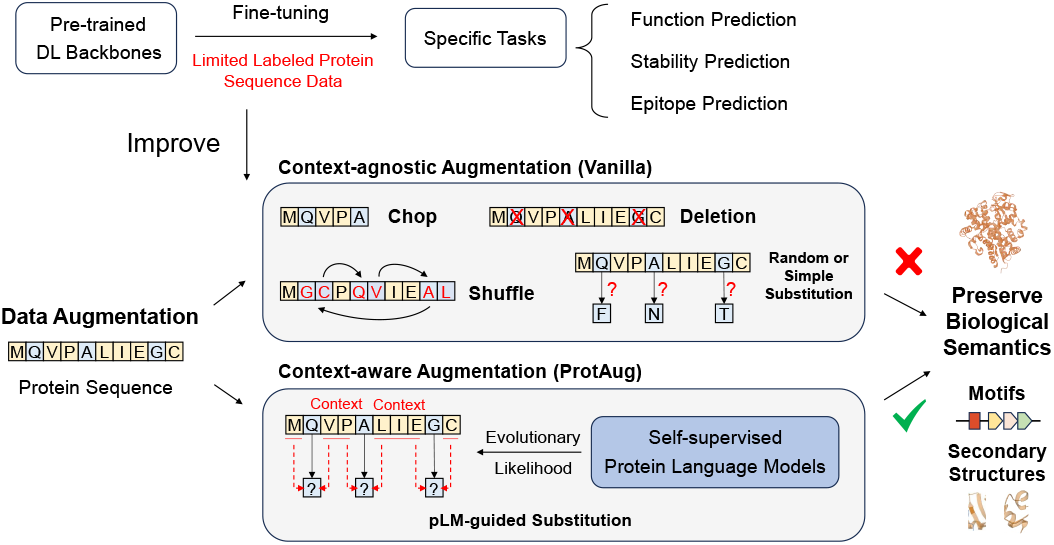
Illustrations of data augmentation on protein sequences. Methods that do not consider protein context tend to disrupt biological semantics, whereas our proposed ProtAug framework utilizes protein language contexts to guide substitutions, thereby better preserving semantics.

Protein language models learn the contextual relationships within protein sequences. During self-supervised pretraining, they distill extensive sequence information into implicit priors that help distinguish between plausible and implausible configurations. Therefore, in this paper, we propose **ProtAug**, a framework that utilizes the contextual information embedded in pLMs, to generate biologically plausible sequences for effective protein data augmentation. Within this framework, we develop two substitution-based methods, namely **ProtAug Esm** and **ProtAug GPT**. The former is guided by the autoencoding model Esm2 [Lin *et al*., 2023a], and the latter uses the autoregressive model ProtGPT2 [Ferruz *et al*., 2022].

Specifically, our proposed methods begin by randomly masking a portion of residues and then resample the masked residues using either a bidirectional or an autoregressive mechanism. We have developed an iterative masking-replacement procedure to allow users to adjust the variation level (0–77%) for task adaptation. Our focus on substitution strategies ensures better preservation of protein semantics, whereas deletion, repetition, and shuffling strategies are commonly believed to disrupt protein structures and functions. Furthermore, ProtAug does not require querying external evolutionary statistics, since this information is already encoded in the pLM parameters.

Experiments conducted on seven tasks using three pLMs address a core question: can the evolutionary likelihood constraints learned by pLMs—typically used for *de novo* generation—be repurposed as a controlled substitution mechanism for data augmentation? To investigate this, we employ a two-tier evaluation.

First, because direct label verification is infeasible at scale, we use generative quality metrics (motif retention, structural similarity, embedding distribution) as proxies for semantic integrity. These metrics are not intended to claim biological novelty or functional validity of the generated sequences; rather, they serve as diagnostic tools to quantify how much original semantic information is preserved. They may help explain why certain augmentation strategies outperform others under specific conditions (e.g., variation level, task type), without assuming that preservation of these metrics guarantees downstream gains.

Second, we measure downstream task performance under controlled conditions (including low-resource settings and duplicate baselines) as the ultimate criterion for augmentation effectiveness. Results show that ProtAug Esm achieves top-two performance in five out of seven tasks at 10% variation, with substantial gains in low-resource (5–20%) data regimes. Further ablation analyses reveal that optimal variation levels are task-dependent: 2–30% works best overall, but higher variation (55–77%) can benefit structure-related tasks, likely via regularization. Semantic preservation correlates with performance at low variation levels, but high-variation augmentation can improve generalization even when label consistency is partially compromised. Based on these findings, we provide task-adaptive recommendations for selecting augmentation methods and variation levels.

### Research Questions and Contributions

This study is guided by four research questions, which collectively aim to provide a systematic empirical understanding of pLM-guided protein sequence augmentation:

- **(Q1) Signal preservation:** Do sequences generated by pLM-guided methods preserve more original biological signals (motifs, structure, and embedding distribution) than simpler alternatives?
- **(Q2) Downstream task effectiveness:** To what extent does augmentation improve prediction performance, particularly in low-resource settings?
- **(Q3) Variation level effects:** How do different variation levels (0–77%) affect downstream task accuracy across augmentation methods, pLM backbones, and task types?
- **(Q4) Necessity of biological plausibility:** Is biological plausibility (i.e., preserving original signals) a necessary condition for achieving downstream improvement, or can regularization-based gains occur even when plausibility is compromised?

## 2 Related Work

### 2.1 Data Augmentation on Images and Texts

Traditional methods for augmenting image datasets include geometric transformations such as rotation, scaling, translation, and flipping, which preserve image characteristics while introducing variability [Krizhevsky *et al*., 2017]. Recent advancements have introduced generative models, such as Generative Adversarial Networks (GANs), to generate realistic synthetic images for augmentation [Goodfellow *et al*., 2014; Gatys *et al*., 2016]. In contrast, text data augmentation methods often involve synonym replacement, replacing words in a sentence with their synonyms to generate semantically similar sentences [Wei and Zou, 2019]. Back-translation, which involves translating text into another language and then back to the original, is also effective at producing diverse textual variations [Sennrich *et al*., 2016]. Advanced approaches leverage LLMs to generate contextually relevant sentences from existing text [Kobayashi, 2018; Dai *et al*., 2025]. Additionally, techniques such as word dropout and random insertion have been explored to introduce noise into the training data, thereby further enhancing model robustness [Wei and Zou, 2019]. In our work, we treat protein sequences as sentences and amino acids as words or tokens, and utilize pLMs rather than LLMs to generate semantics-preserving proteins based on the context surrounding the positions to be edited.

### 2.2 Data Augmentation on Protein Sequences

There is limited work on exploring augmentation methods for protein sequences compared to images and texts. The challenges lie in the unique characteristics of protein sequences compared to natural language and image: (1) protein sequences have a small vocabulary of only 20 amino acids and a few special tokens, making every edit in the sequence a risk of causing substantial changes in structure and function and (2) it is difficult to verify that a transformation preserves the original label of a sequence, whereas the fidelity of an augmented sentence or image can be inspected handily. Current methods circumvent the semantic-checking issue using external rules. For example, *Retrieved Sequence Augmentation* [Ma *et al*., 2023] retrieves existing homologous sequences from large databases, which is time-consuming. In contrast, PAM matrices [Oliveira *et al*., 2023] provide evolutionary frequencies for substitution but ignore contextual information. Domain-motivated transformation has been found to achieve consistent results in contrastive learning [Shen *et al*., 2021], but requires manual editing based on a priori motifs.

Random substitution and other simple strategies, such as repetition, deletion, and shuffling, are explored but are likely to disrupt semantics [Sun *et al*., 2024]. In a recent comprehensive benchmark, Sun et al. [2024] extended image/text augmentation techniques to proteins and proposed two semantic-level augmentation methods: Integrated Gradients Substitution and Back Translation Substitution. While their work represents an important advance, we encountered reproducibility issues when attempting to run their code (see Section 4.3). Moreover, their evaluation focused on the Automated Protein Augmentation (APA) framework that adaptively selects augmentation combinations, leaving open the question of how individual pLM-guided strategies compare to simpler baselines under controlled conditions.

Gibbs sampling, which iteratively redraws amino acids using the probabilities predicted by transformer models, has been used to generate sequences [Rao *et al*., 2020] but has not been tested as an augmentation method. In this paper, we propose an augmentation framework that uses the evolutionary prior already distilled inside the pre-trained pLM to decide which substitutions are biologically plausible without any external look-up or manual masking, and we further assess the fidelity of the augmented sequences.

## 3 Methods

### 3.1 Problem Formulation

The purpose of our proposed ProtAug framework is semantic-level augmentation for protein sequences. When the number of labeled protein sequences in a dataset is insufficient, we use ProtAug to generate novel sequences from existing sequences for accuracy improvement by introducing variation while preserving the properties of the original data.

In this paper, we define the protein augmentation problem as follows: given an original protein sequence

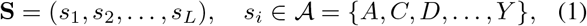

where *L* is the length of the sequence and *s*_*i*_ is one of the 20 amino acids *A*, generate an augmented sequence *S*^*′*^ with the normalized Hamming distance (denoted as **variation level**) defined as

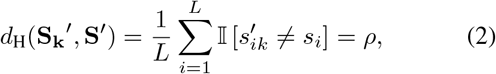

for a user-specified variation level *ρ* ∈ [0, 1], while preserving the associated biological label of **S**. *k* is the number of iterations required to reach the specified variation level *ρ*.

### 3.2 ProtAug Esm: Encoder-pLM-guided Substitution

Current pLMs are of the transformer architecture, consisting of encoders, decoders, or both [Elnaggar *et al*., 2021]. As a pLM with both encoders and decoders works similarly to a decoder-only pLM for generating protein sequences, in this paper we study two methods in ProtAug: one with the encoder and the other with the decoder.

Models adopting the encoder architecture are good at providing contextual information across the entire sequence and can infer a portion of masked positions based on the amino acids preceding or following them. Models in the Prot-Trans program, such as ProtBert [Elnaggar *et al*., 2021] and Esm2 [Lin *et al*., 2023a], are in this category.

In this study, we use Esm2 for the encoder-based strategy, given Esm2’s superior performance on several downstream tasks. As illustrated in Figure 2A and Algorithm 1, to generate a new sequence from a given one, we first randomly mask a portion of the amino acids in the sequence. From preliminary experiments, we find that Esm2 yields many repetitive amino acids when masking a large portion of the data. To alleviate this issue, we restrict the masked portion to a maximum of 0.15, which is the default value for pretraining BERT. Starting from the masked position, 80% remain masked, 10% are replaced with a random amino acid, and the remaining 10% are restored to their original residues, following the pretraining procedure in the BERT masked language modeling task [Devlin *et al*., 2019]. In Step 2, Esm2 is used to generate logits, which are further converted to probabilities, and then amino acids are redrawn using these probabilities. When a generated sequence has a variation level higher than 0.15, we perform multiple rounds of sampling until the variation level is reduced to the threshold. If a sequence is longer than 1000 tokens (amino acids), we split it into slices of 1000 tokens and process each slice in turn.

**Figure 2.**
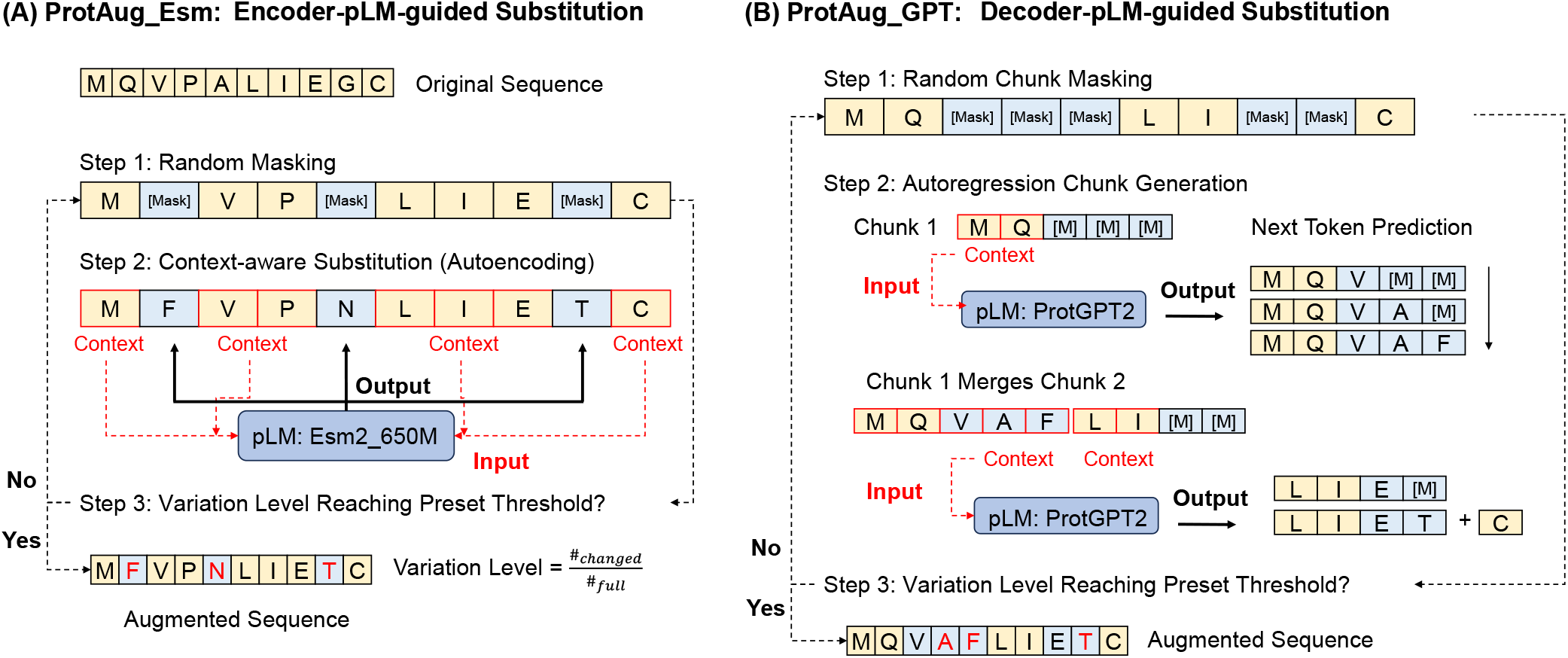
Illustrations of the two semantics-preserving substitution augmentation methods in ProtAug. In both methods, a portion of the amino acids is randomly masked, and then protein language models are utilized to predict the amino acids in these masked positions. ProtAug Esm uses Esm to generate all amino acids at once by an autoencoding mechanism, whereas ProtAug GPT uses ProtGPT2 to generate next tokens chunk by chunk. Both methods adopt an iterative procedure for the variation level to reach the user-specified threshold.

### 3.3 ProtAug GPT: Decoder-pLM-guided Substitution

Models adopting the decoder architecture generate new tokens either from a given fragment or from scratch in a leftto-right manner. ProtGPT2 is a generative model for protein sequences in this category. Because the decoder model generates new tokens based on the preceding (or left) context, uniformly masking random positions requires tens or hundreds of passes to fully generate a new protein sequence of the same length.

To speed up generation, we group masked positions into chunks. Specifically, when the variation level is less than 0.1, we generate 3 chunks; when it is between 0.1 and 0.4, we generate 5 chunks; for levels above 0.4, we generate 10 chunks (see Figure 2B and Algorithm 2).

In Step 1, we split each slice of the protein sequence into the predefined number of chunks *c* and randomly sample a starting position for each chunk. The total number of masked positions in each slice is calculated as the variation level *ρ* multiplied by the length of the slice. Then we divide this number by the number of chunks to obtain the average number of masked positions *w* for each chunk in a slice. From the sampled starting position in each chunk, we mask the following amino acids from left to right until we reach the specified number of masked positions for that chunk or reach another masked position. Consecutive masked positions across chunks are merged as one chunk, further accelerating the inference pass of ProtGPT2. After merging, a chunk is a fragment with consecutive masking positions at its right end.

#### Algorithm 1 ProtAug Esm: Encoder-pLM-guided Substitution

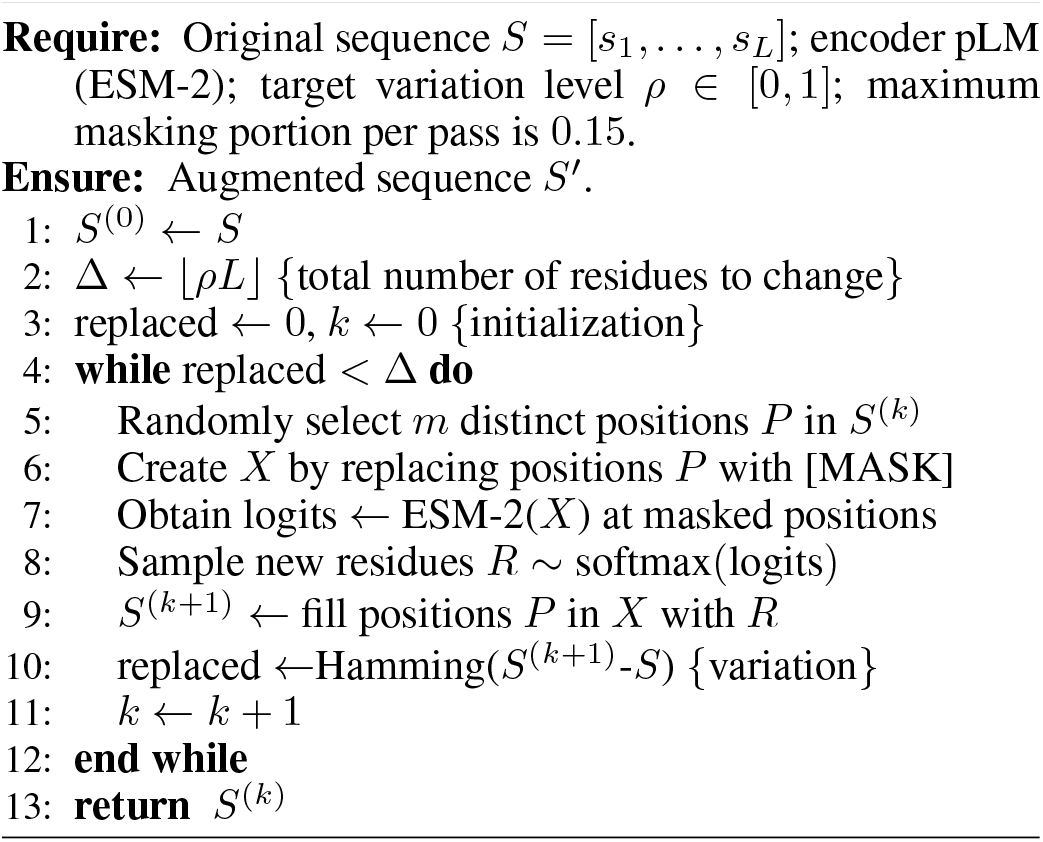

In Step 2, we use the masked chunk 1 sequence as input to the ProtGPT2 model to generate amino acids at the masked positions. For all the ‘\ n’ tokens generated by ProtGPT2, we replace them with the neutral and frequent amino acid ‘L’ (Leucine). If ProtGPT2 does not return a sequence of the required length, we supplement the remainder with random amino acids. After all masked positions are filled, chunk 1 is further merged with the masked chunk 2 to form a larger masked sequence, which is then used as input for ProtGPT2. This merge-and-predict loop continues until all masked positions in all chunks are filled, and then the variation level of the generated sequence is checked. We perform multiple rounds of sampling until the variation level reaches the threshold. Since ProtGPT2 can only handle sequences of fewer than 1280 tokens, we split each sequence into slices of 1000 tokens.

## 4 Experiments and Results

### 4.1 Experimental Settings

#### Datasets and Workloads

We use five datasets, which give seven prediction tasks, to evaluate the effectiveness of our ProtAug framework. The first three datasets are sourced from Kaggle competitions and the other four are collected from the TAPE benchmark [Rao *et al*., 2019]: (1) **Protein function prediction** on the Molecular Function aspect **(Function MF)** involves predicting the presence of 677 functions defined by Gene Ontology [Aleksander *et al*., 2023]. (2) **Thermostability prediction** is to predict the melting temperature of natural enzyme sequences and engineered proteins. (3) **Epitope prediction** provides peptide records which contain the protein sequences to which they belong, the start and end positions within the corresponding proteins, and binary labels indicating whether they are epitopes. (4) **Remote homology** detection on **Family** level classifies proteins into 3888 different structural categories. (5) **Super-family** level homology classifies proteins into 1961 structural categories. (6) **Fold** level homology classifies proteins into 1195 structural categories. (7) **Secondary structure prediction** is to predict the type of local conformation of each amino acid in a protein, with categorical labels *y* ∈ { 0, 1, 2 }. For epitope prediction, we randomly split proteins into training, validation, and test sets at a ratio of 7:2:1. For other tasks, we use the default dataset splits provided in the competitions or in TAPE. We summarize the statistics of the five datasets used in this study in Appendix A.

##### Algorithm 2 ProtAug GPT: Decoder-pLM-guided Substitution

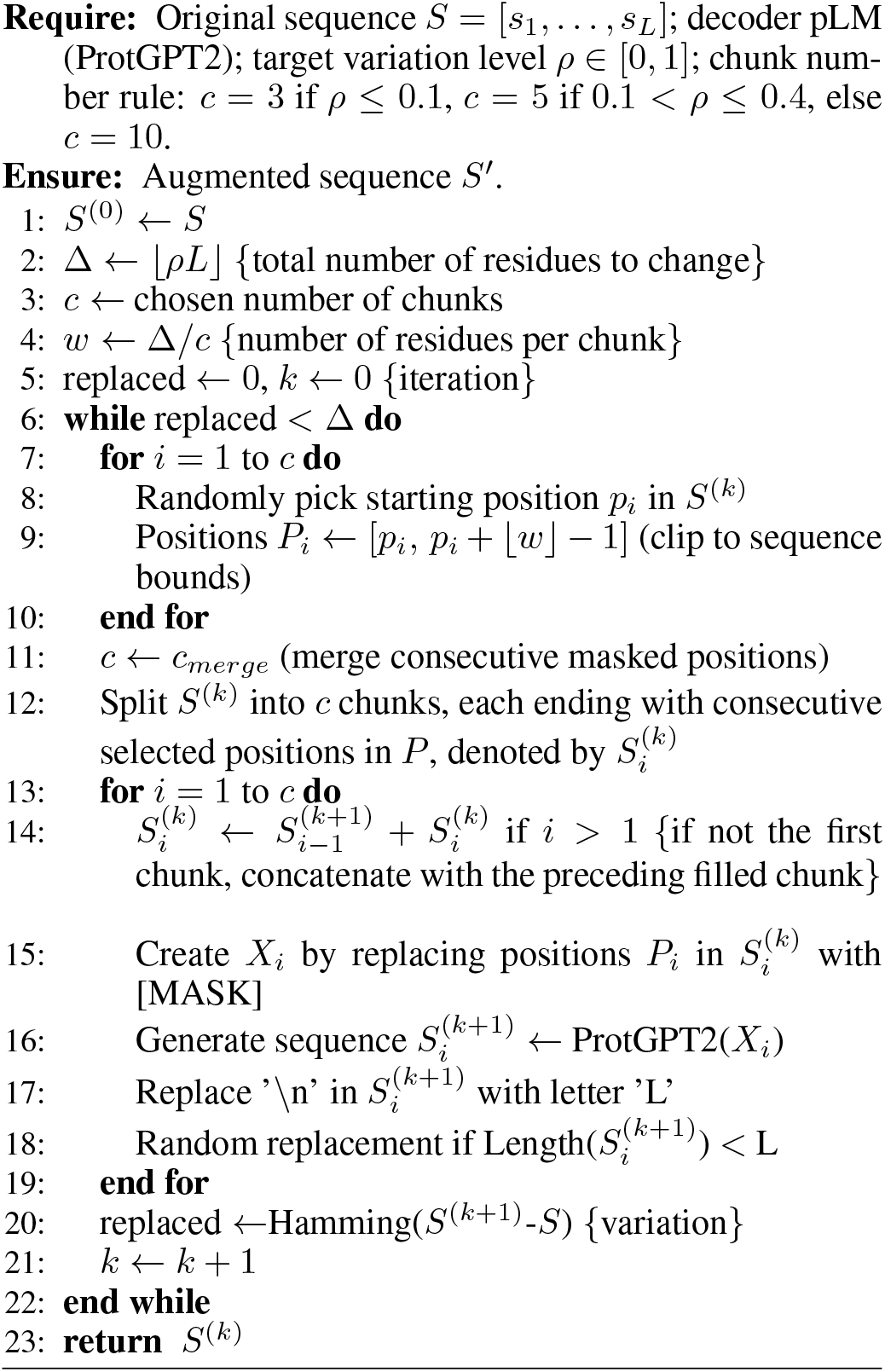

For function prediction, we use Fmax score as the evaluation metric, following the CAFA competition [Zhou *et al*., 2019] settings. For stability prediction, we calculate the Spearman correlation coefficient **(rho)** between the predicted and ground truth values for evaluation. For epitope prediction, we adopt the Area Under the Curve **(AUC)**, which measures the area under the ROC (Receiver Operating Characteristic) curve, as the evaluation metric. For other tasks, the vanilla accuracy is used.

#### Platform

All experiments are conducted on a server with an NVIDIA RTX 3090 GPU with 24 GB GPU memory. The ProtAug framework is implemented in TensorFlow 2.0 [Abadi *et al*., 2016]. We have modified the ktrain [Maiya, 2020] source code to support the Esm2 model, as Esm2 is not supported in the original ktrain.

#### Fine-tuning

The backbones of pLMs are fixed and only prediction heads are trained. All models are fine-tuned with early stopping if the validation loss is greater than that from the previous epoch. We also explored full-parameter fine-tuning but found it prone to overfitting; therefore, we report only fixed-backbone training in this study. More details and results are reported in Appendix B and D.

### 4.2 Metrics for Sequence Quality Evaluation

#### Variation Level

We use the normalized Hamming distance to measure residue-wise amino acid differences between the original and the generated sequences. We first count positions in the two sequences that differ in residues, then divide this count by the length of the sequence.

#### Motif Retention Rate

We perform a homology search with *hmmscan* across the Pfam database [Paysan-Lafosse *et al*., 2025] and retain aligned motifs with E-values greater than 1e-4. The **motif retention rate** is calculated as the percentage of overlapping motifs in the original motifs. We also count the number of disrupted proteins caused by substitution. A protein is considered disrupted if no Pfam motif is aligned with it.

#### Amino Acid Diversity

We use Shannon diversity to measure amino acid diversity in a sequence based on the occurrences of amino acids (Appendix C).

#### Amino Acid Repetition

The repetition ratio of amino acids in a sequence is defined as the number of subsequences of three identical amino acids divided by the length of the sequence. For example, the repetition ratio of MDPAAAA is 2/7 because the sequence is of length 7 and contains two AAA’s (Appendix C).

#### Structure Analysis

In order to assess whether a generated protein retains the structural motif of the original protein, we use EsmFold to predict the structure of a protein from its amino acid sequence. Due to the limitation of computational resources, we perform prediction only for proteins less than 713 aa.. We use TM-score [Zhang and Skolnick, 2005] to quantify the similarity between two structures, and use pLDDT from EsmFold [Lin *et al*., 2023b] to indicate the quality of the prediction. Higher values of TM-score and pLDDT indicate greater similarity between the predicted structure of the generated protein and the original protein, and better prediction quality, respectively. Structures of original proteins are retrieved from AlphaFoldDB [Varadi *et al*., 2022].

### 4.3 Baseline Methods

#### No Augmentation

**No aug** denotes not using any augmented data. Without augmentation, the number of training samples is about half of those with augmentation.

#### Duplicate

We duplicate the training data as a control for sample size in the training set when investigating the effect of different variation levels (denote as **Duplicate**).

#### Random Substitution

A portion of the amino acids in a sequence is randomly masked, and each masked position is randomly replaced by one of the 20 amino acids (denoted as **Random**).

#### PAM-guided Substitution

Instead of fully random sampling, different PAM (Point Accepted Mutation) matrices [Dayhoff *et al*., 1978] are used to provide evolutionary hints for substitution (denoted as **PAM**). Each element in a PAM matrix represents the probability of one amino acid being replaced by another. Lower-numbered PAM matrices correspond to more conservative substitutions, while higher-numbered matrices allow for greater variation. We utilize various PAM matrices to generate sequences with specified levels of variation: PAM1 for 2% and 10%, PAM50 for 30%, and PAM250 for 55% and 77%.

#### Homology Search

The SwissProt [Uniprot, 2023] database, which contains around 570,000 natural sequences, is used as the external resource. After filtering database sequences that share similarities greater than 90% with sequences in training, validation, and test sets, we search each protein in training sets against SwissProt with *Diamond* and use the sequence with the highest identity as replacement for augmentation. Results with E-value less than 1e-10 and identity value less than 0.2 are filtered. Different variation levels are set as identity upper bounds, and sequences with no hits are replaced with generated sequences in **PAM** with the same variation level.

### 4.4 Q1: Signal Preservation — Do pLM-guided Methods Retain More Original Information?

We measure the quality of generated sequences by motif retention, amino acid diversity, repetition, and structure similarity. **HomologyTop** denotes that we set identity **lower bound** instead of upper bound as in Homology, to retrieve the most similar sequences in SwissProt. These two homology search methods, as they contain natural proteins, are served as positive control. Overall, ProtAug Esm preserves motifs, structure, and embedding distribution better than Random/PAM, comparable to Homology.

In Figure 3, with increasing variation level, protein motifs are more widely disrupted, and more proteins lose their original motifs completely, as reflected by decreasing motif retention rates and increasing numbers of disrupted proteins. Our two proposed pLM-guided methods, especially ProtAug Esm, retain original semantic signals comparable to natural sequences from homology search, and better than the two simple substitution methods, reflected by their higher motif retention rates and lower disruption counts. This could partly explain why they show advantages over context-agnostic substitution methods such as Random and PAM-guided substitution.

**Figure 3.**
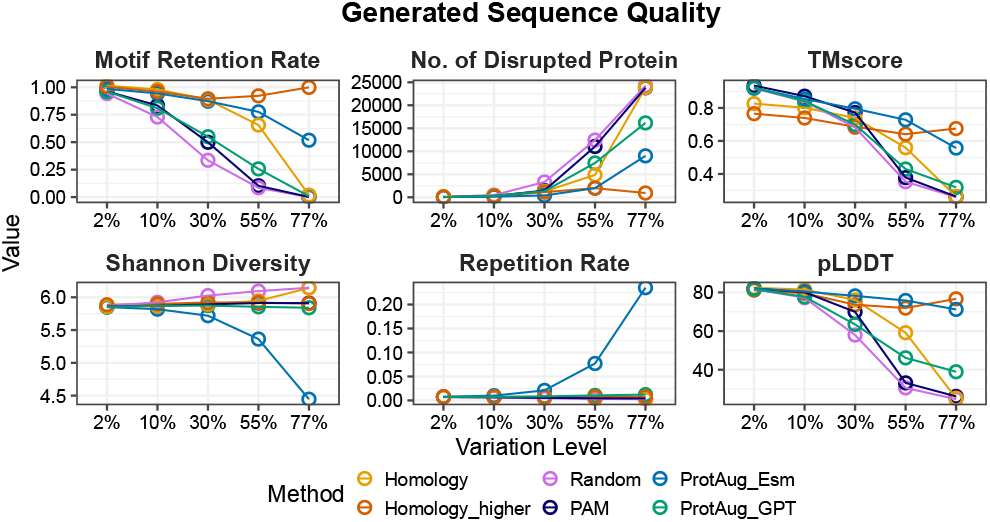
Quality of the augmented data on the Function MF dataset.

We also observe that the Shannon diversity decreases and the repetition ratio increases dramatically when using ProtAug Esm but not so when using ProtAug GPT. This could be due to the bi-directional autoencoding mechanism in which amino acids at masked positions are generated simultaneously. This mechanism prioritizes amino acids with high occurrences in the training data when the original protein context is not sufficiently preserved. In comparison, Pro-tAug GPT preserves high amino acid diversity, which may be beneficial for specific tasks.

Furthermore, we use TM-score to measure the structural similarity between the augmented and the original sequence, and use pLDDT to measure the confidence of structure prediction. We find that sequences generated by ProtAug Esm are more similar to their original ones in structures compared to other methods, even more similar than sequences from homology search. Predicted structures from ProtAug-generated sequences have confidence comparable to that of Homology-Top natural sequences and higher than that of other augmentation methods.

The t-SNE visualization in Figure 4 demonstrates that pLM-generated proteins are more similar to the original ones than those generated by simple methods, as indicated by smaller separation between the two clusters in ProtAug Esm embeddings, especially at higher variation levels (*>*30%).

**Figure 4.**
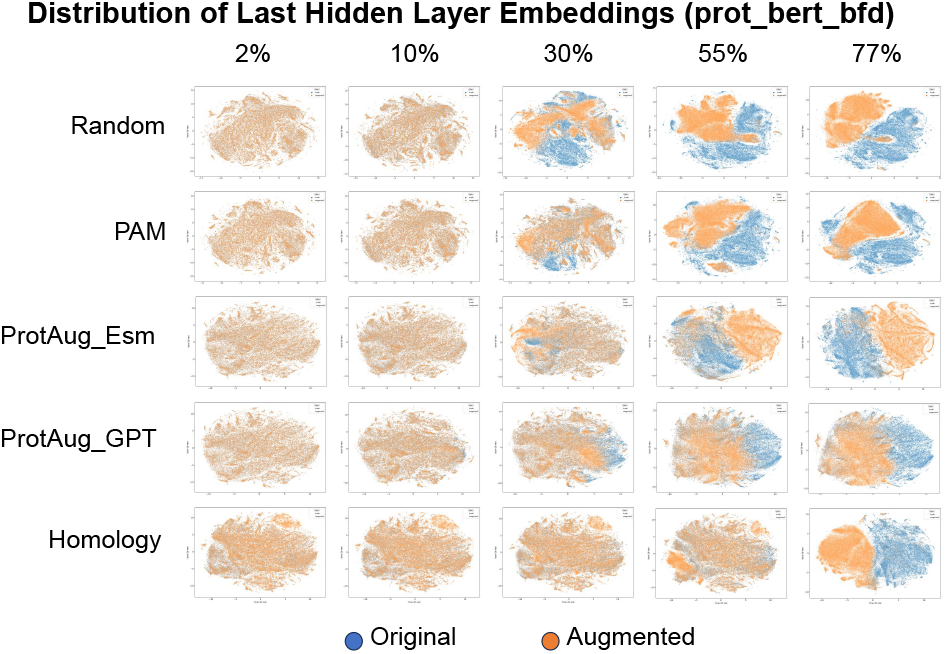
Distribution drifts between the original and generated data in Function MF dataset shown by t-SNE plot.

### 4.5 Q2: Downstream Task Effectiveness — How Much Improvement Is Achieved, Especially in Low-Resource Settings?

We evaluate different augmentation strategies with a 1:1 generation at a 10% variation level for fine-tuning three widely used pLMs in seven tasks.

Table 1 shows that ProtAug Esm achieves top-2 performance in 5 out of 7 tasks (4 rank-1), and ProtAug GPT, while less effective than ProtAug Esm, wins in 3 out of 7 tasks (3 rank-2). Substitution augmentation generally improves prediction accuracy compared to the No aug baseline and even over Homology, which uses external resources. Among the three pLM backbones, Esm2 generally produces the best results. These findings highlight the robustness and effectiveness of our proposed ProtAug framework, especially ProtAug Esm, in fine-tuning pLMs for performance improvement. We further demonstrate that ProtAug Esm performs reasonably well in low-resource settings, across five smaller fractions ranged from 1% to 50% training data in all seven tasks, achieving the best performance in most cases over other augmentation methods. (Figure 5; also see Appendix E).

**Table 1.** Performance of different augmentation across pLMs and tasks with mean values and standard deviations. Values in bold and underlined indicate the highest and second-highest performance, respectively, for a specific prediction model and task. Shaded cells indicate the highest and second-highest performance in a certain task across all settings.

| Model<br>(Parameters) | Method | Function MF | Stability | Epitope | Family | Remote Homology<br>Superfamily | Fold | Secondary<br>Structure |
| --- | --- | --- | --- | --- | --- | --- | --- | --- |
| DistilProtBert (230M) | No_aug | 0.562 ($\pm 0.008$) | 0.494 ($\pm 0.006$) | 0.514 ($\pm 0.009$) | 0.216 ($\pm 0.002$) | 0.035 ($\pm 0.003$) | 0.087 ($\pm 0.013$) | 0.777 ($\pm 0.003$) |
| | Homology | **0.579 ($\pm 0.002$)** | 0.495 ($\pm 0.010$) | 0.494 ($\pm 0.032$) | 0.271 ($\pm 0.022$) | 0.041 ($\pm 0.009$) | 0.081 ($\pm 0.007$) | 0.771 ($\pm 0.002$) |
| | Random | 0.578 ($\pm 0.003$) | 0.512 ($\pm 0.014$) | 0.470 ($\pm 0.019$) | 0.340 ($\pm 0.011$) | **0.051 ($\pm 0.006$)** | 0.087 ($\pm 0.008$) | 0.779 ($\pm 0.002$) |
| | PAM | 0.578 ($\pm 0.004$) | **0.517 ($\pm 0.015$)** | 0.501 ($\pm 0.025$) | 0.352 ($\pm 0.008$) | 0.047 ($\pm 0.012$) | 0.088 ($\pm 0.009$) | 0.778 ($\pm 0.004$) |
| | ProtAug_Esm | 0.573 ($\pm 0.001$) | 0.511 ($\pm 0.014$) | **0.539 ($\pm 0.011$)** | **0.363 ($\pm 0.012$)** | 0.051 ($\pm 0.007$) | 0.084 ($\pm 0.011$) | 0.777 ($\pm 0.007$) |
| | ProtAug_GPT | 0.573 ($\pm 0.004$) | 0.510 ($\pm 0.026$) | 0.510 ($\pm 0.036$) | 0.331 ($\pm 0.008$) | 0.051 ($\pm 0.004$) | **0.091 ($\pm 0.008$)** | **0.780 ($\pm 0.002$)** |
| prot_bert_bfd (430M) | No_aug | 0.620 ($\pm 0.005$) | 0.610 ($\pm 0.006$) | 0.619 ($\pm 0.027$) | 0.271 ($\pm 0.009$) | 0.073 ($\pm 0.002$) | **0.170 ($\pm 0.020$)** | 0.818 ($\pm 0.001$) |
| | Homology | **0.624 ($\pm 0.002$)** | 0.577 ($\pm 0.007$) | **0.662 ($\pm 0.019$)** | 0.406 ($\pm 0.017$) | 0.096 ($\pm 0.002$) | 0.164 ($\pm 0.011$) | 0.813 ($\pm 0.002$) |
| | Random | 0.620 ($\pm 0.004$) | 0.598 ($\pm 0.002$) | 0.600 ($\pm 0.015$) | 0.487 ($\pm 0.013$) | 0.112 ($\pm 0.002$) | 0.165 ($\pm 0.016$) | 0.819 ($\pm 0.002$) |
| | PAM | 0.615 ($\pm 0.004$) | **0.616 ($\pm 0.006$)** | 0.632 ($\pm 0.010$) | 0.497 ($\pm 0.014$) | 0.117 ($\pm 0.010$) | 0.166 ($\pm 0.018$) | **0.820 ($\pm 0.002$)** |
| | ProtAug_Esm | 0.620 ($\pm 0.001$) | 0.597 ($\pm 0.009$) | **0.677 ($\pm 0.010$)** | **0.504 ($\pm 0.013$)** | **0.120 ($\pm 0.008$)** | 0.169 ($\pm 0.015$) | 0.819 ($\pm 0.001$) |
| | ProtAug_GPT | 0.617 ($\pm 0.003$) | 0.610 ($\pm 0.007$) | 0.634 ($\pm 0.008$) | 0.487 ($\pm 0.009$) | 0.109 ($\pm 0.010$) | 0.166 ($\pm 0.020$) | 0.819 ($\pm 0.002$) |
| Esm2 (650M) | No_aug | 0.659 ($\pm 0.005$) | 0.641 ($\pm 0.019$) | 0.470 ($\pm 0.009$) | 0.481 ($\pm 0.006$) | 0.305 ($\pm 0.142$) | **0.284 ($\pm 0.007$)** | **0.825 ($\pm 0.002$)** |
| | Homology | 0.661 ($\pm 0.007$) | 0.609 ($\pm 0.012$) | 0.455 ($\pm 0.006$) | 0.767 ($\pm 0.016$) | 0.418 ($\pm 0.002$) | 0.278 ($\pm 0.005$) | 0.806 ($\pm 0.003$) |
| | Random | 0.658 ($\pm 0.008$) | 0.640 ($\pm 0.002$) | 0.475 ($\pm 0.012$) | 0.845 ($\pm 0.002$) | **0.476 ($\pm 0.002$)** | 0.276 ($\pm 0.010$) | 0.822 ($\pm 0.003$) |
| | PAM | 0.656 ($\pm 0.003$) | **0.647 ($\pm 0.002$)** | 0.470 ($\pm 0.017$) | **0.848 ($\pm 0.002$)** | 0.470 ($\pm 0.007$) | 0.275 ($\pm 0.013$) | 0.825 ($\pm 0.004$) |
| | ProtAug_Esm | **0.663 ($\pm 0.003$)** | 0.644 ($\pm 0.005$) | **0.514 ($\pm 0.022$)** | **0.850 ($\pm 0.005$)** | 0.471 ($\pm 0.004$) | 0.278 ($\pm 0.010$) | **0.831 ($\pm 0.004$)** |
| | ProtAug_GPT | 0.661 ($\pm 0.006$) | **0.646 ($\pm 0.002$)** | 0.482 ($\pm 0.013$) | 0.836 ($\pm 0.010$) | 0.455 ($\pm 0.003$) | **0.280 ($\pm 0.011$)** | 0.823 ($\pm 0.004$) |

**Figure 5.**
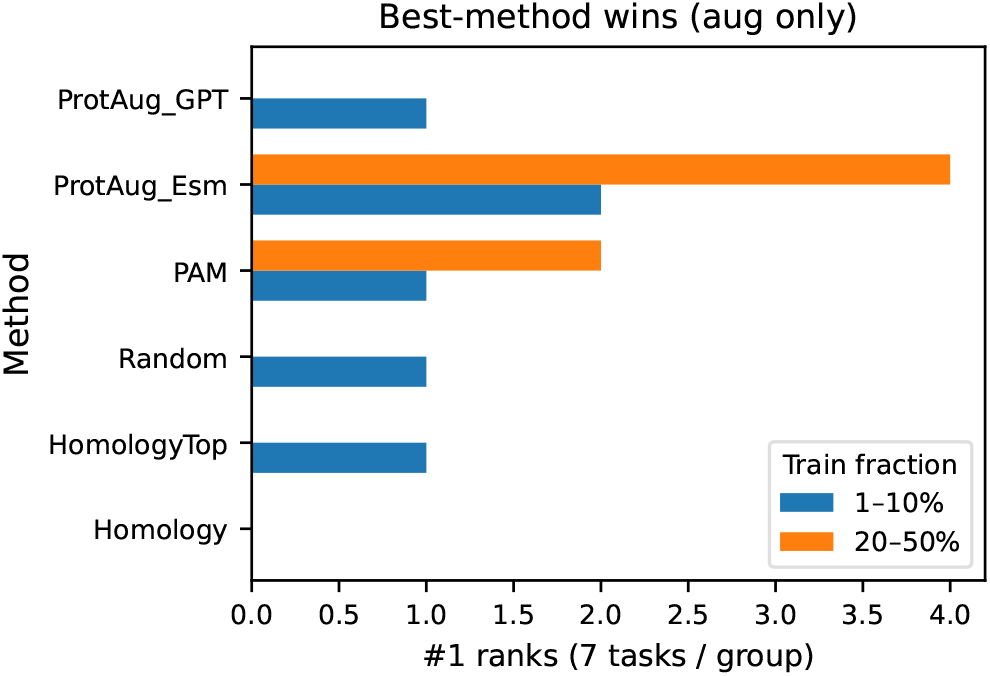
Number of times each augmentation method achieved the best performance across 7 tasks and 5 low-resource data fractions (1%, 5%, 10%, 20%, 50%) using Esm2 backbone.

### 4.6 Q3: Variation Level Effects — How Does Substitution Rate Affect Performance?

We also investigate the performance effects of using augmentation data generated from different methods with 0%, 2%, 10%, 30%, 55%, and 77% differences (variation levels) from their original sequences. For a fair comparison, we define 0% as using duplicate (denoted as **Duplicate**) training data to ensure equal training set sizes across all variation levels.

Results indicate that the effect of variation level differs among prediction models, augmentation methods, and tasks (Figure 6). We report results of three tasks and the complete results can be found in Appendix F. We observe that augmentations with 2% and 10% variation levels tend to outperform the No aug setting, with 10% variation achieving the overall best performance. Surprisingly, using homology search with the most similar sequences (HomologyTop) as augmentation does not guarantee the best results, potentially due to label inconsistency. In general, the best performance falls whitin a 2% to 30% variation range. However, we observe that Pro-tAug GPT with a variation level of 77% achieves high performance gain on secondary structure prediction. The two smaller models also benefit from high-variation-level augmentation in some cases. These results suggest that a flexible variation level can be beneficial depending on specific augmentation methods, pLM models, and tasks.

**Figure 6.**
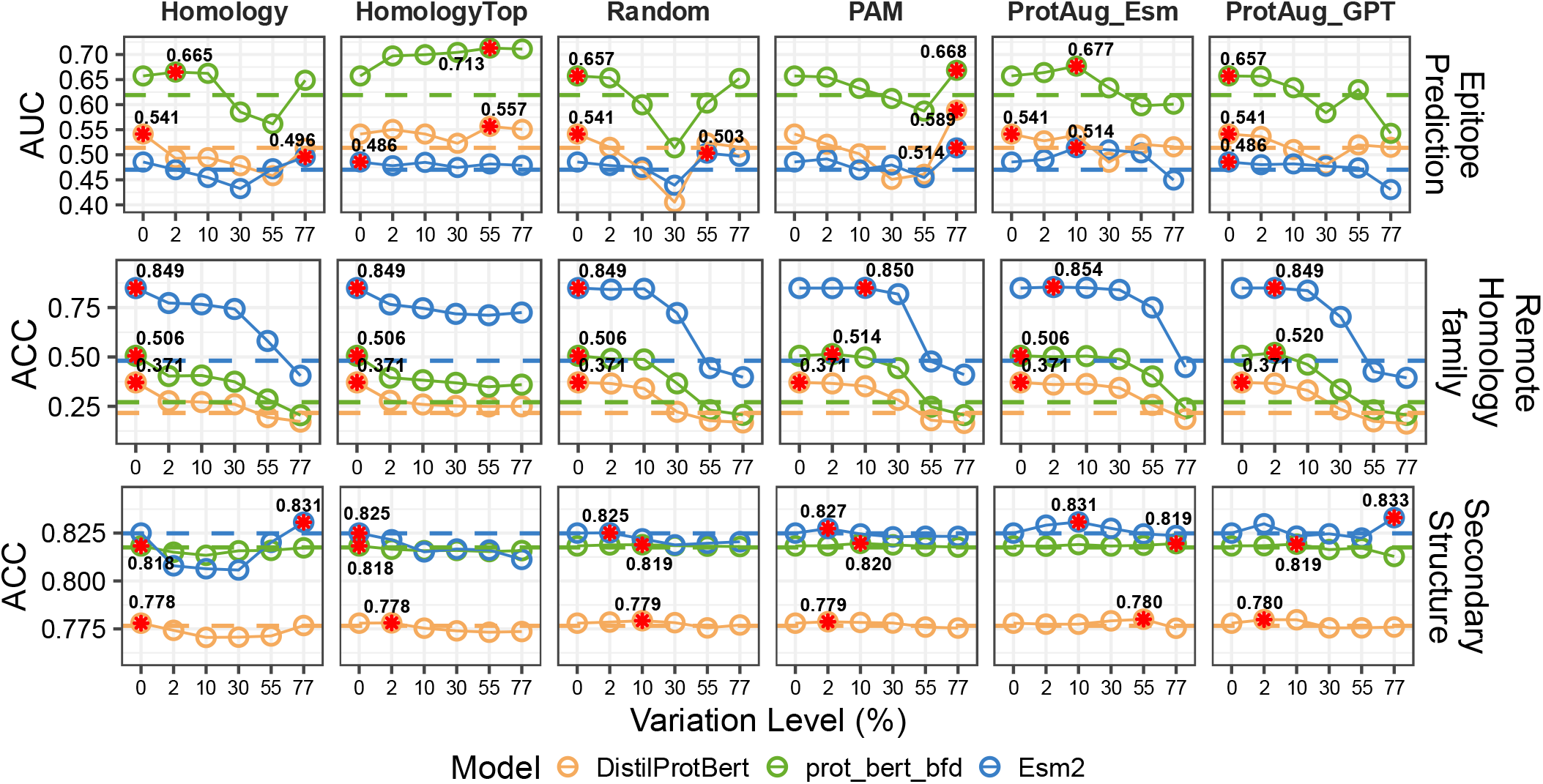
Performance of fine-tuning pLMs across variation levels. Dashed lines indicate performance without augmentation. Stars denote the highest scores of corresponding models.

In addition, we assess the average time required for augmentation for these strategies. In Table 2, the two simple substitution methods are the most time-efficient, whereas the augmentation time of the two pLM-guided methods increases with the variation level due to the iteration of sampling and replacement.

**Table 2.**
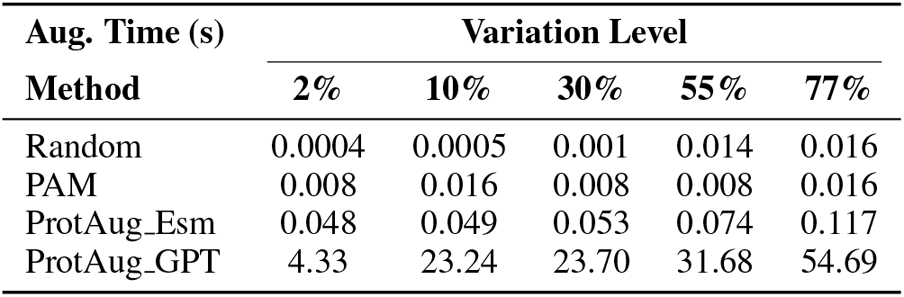
Average augmentation time of protein sequence in Epitope with average length of 572 aa.

Considering both the accuracy performance and augmentation time, we recommend ProtAug Esm over ProtAug GPT at the 2-30% variation level in future studies.

### 4.7 Q4: Necessity of Biological Plausibility — Is Semantic Preservation Required for Improvement?

To assess whether biological plausibility (i.e., preserving original sequence signals) is necessary for achieving downstream improvement, we examine the relationship between signal preservation (Q1) and performance gains (Q2–Q3) across variation levels and task types. To evaluate whether the augmented sequences retain the original labels and help improve learning generalization, we use embeddings of training samples as nearest-neighbor predictors and transfer labels to test samples. The relatively high KNN accuracies of ProtAug in corresponding tasks indicate that (1) the feature space of the training data lies close to that of the test set and (2) the labels in the corresponding neighborhood are consistent with the ground-truth labels.

At 2%-30% variation level, ProtAug Esm preserves motifs, structural similarity, and embedding distribution substantially better than Random or PAM (Figure 3 and 4). KNN label consistency remains high (Figure 7), indicating that augmented sequences largely retain the original labels. Under these conditions, we observe consistent performance improvements across most tasks (Table 1, Figure 6). This pattern supports the label-preserving augmentation hypothesis: when semantic integrity is maintained, augmentation effectively expands the training set with valid label-carrying examples.

**Figure 7.**
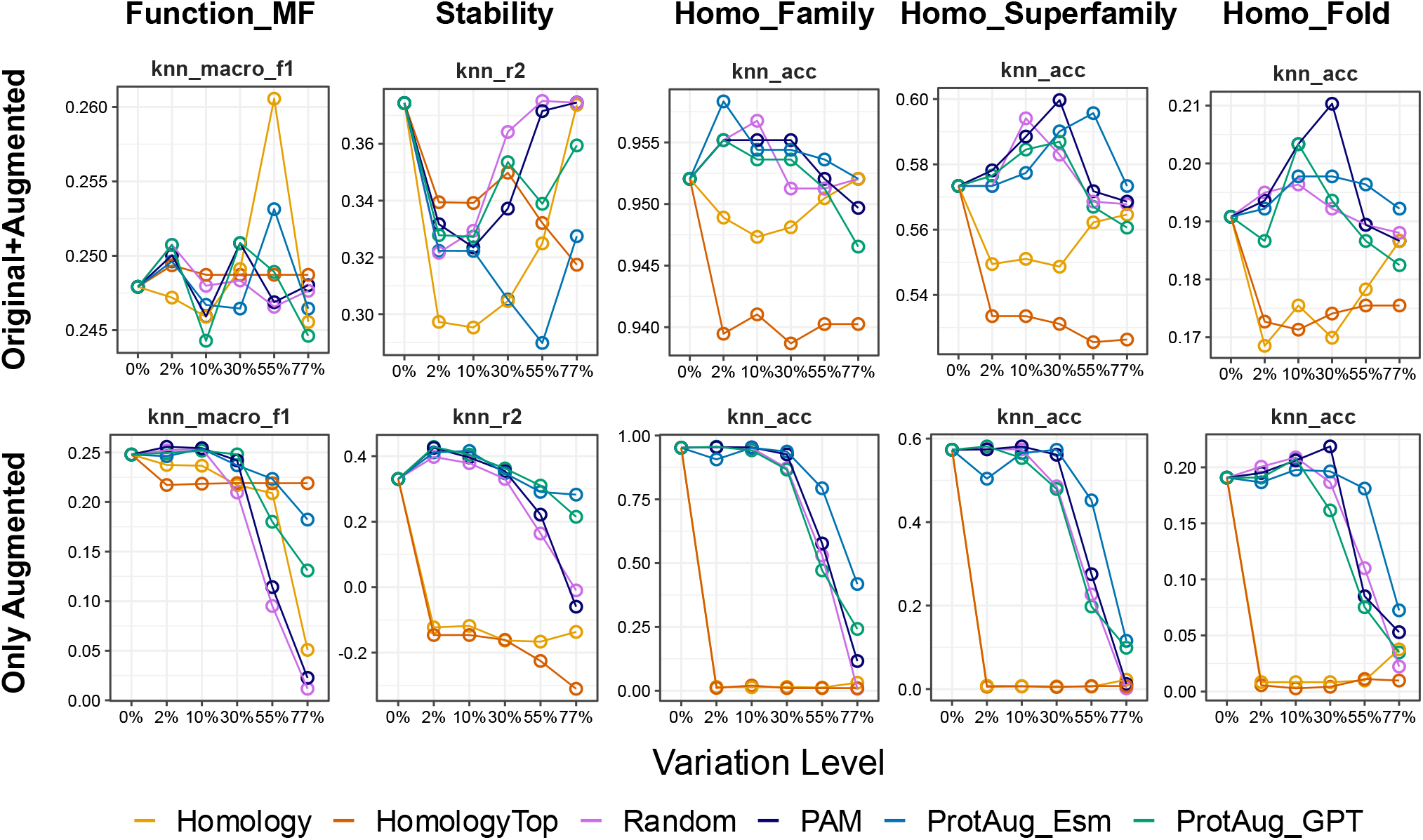
Label consistency measured by KNN predictors.

Interestingly, for structure-rich tasks (superfamily and fold levels of homology detection), KNN label consistency does not decrease—it even increases with higher variation levels. This suggests that structural labels may be more tolerant to sequence variation, challenging the necessity of exact sequence-level plausibility.

At higher levels (55-77%), signal preservation and KNN label consistency decrease substantially. If biological plausibility were strictly necessary, performance should decline correspondingly. However, we observe that ProtAug GPT with 77% variation achieves competitive or even improved performance compared to lower variation levels in secondary structure prediction (Figure 6). This occurs despite clear evidence that many individual residue labels are no longer valid after high-rate substitution, since per-position secondary structure annotations are highly sequence-dependent. It suggests that performance gains at high variation levels arise primarily from regularization rather than from label-preserving sample expansion.

Thus, biological plausibility (semantic preservation) is a sufficient but not necessary condition for effective augmentation. At low-to-moderate variation levels (2–30%), preserving original signals correlates with performance gains, supporting the label-preserving hypothesis. At high variation levels (55–77%), performance can still improve, particularly for structure-related tasks or when overfitting is severe, due to regularization effects.

## 5 Conclusion and Recommendations

In this study, we conducted a systematic empirical investigation of pLM-guided data augmentation for protein sequence prediction tasks. We proposed ProtAug, a framework that leverages both encoder-based (ESM-2) and autoregressive (ProtGPT2) pLMs to generate augmented sequences with controllable variation levels. Our investigation focuses on four questions concerning signal preservation, downstream effectiveness, variation level effects, and the necessity of biological plausibility.

We summarize the answers to the four research questions posed in the Introduction:

- **Q1 (Signal preservation):** ProtAug Esm consistently preserves motifs, structural similarity (TM-score), and embedding distribution (lower MMD^2^) better than Random and PAM, often comparable to Homology retrieval, especially at low-to-moderate variation levels ( ≤ 30%). This supports the claim that pLM-guided methods produce augmented sequences closer to the original data manifold.
- **Q2 (Downstream effectiveness):** Augmentation yields consistent but task-dependent improvements. At 10% variation, ProtAug Esm achieves the best or second-best performance in 5 out of 7 tasks. In low-resource settings (1–50% training data), ProtAug Esm outperforms both No aug and Duplicate with margins up to 0.03–0.05 Fmax, achieving the best performance in most scenarios, underscoring its practical value when labeled data are scarce.
- **Q3 (Variation level effects):** Low-to-moderate variation (2–30%) performs best overall across tasks and models. However, high-variation augmentation (55–77%) can benefit specific tasks (e.g., secondary structure prediction with ProtAug GPT), and smaller pLM backbones occasionally benefit from higher variation. This task- and model-dependent behavior suggests that variation level should be treated as a tunable hyper-parameter.
- **Q4 (Necessity of biological plausibility):** The necessity of plausibility is task- and variation-dependent. At ≤30% variation, higher motif retention and structural similarity correlate with better downstream performance, consistent with the label-preserving augmentation hypothesis. At higher variation levels, especially with ProtAug GPT, performance can improve despite reduced label consistency, suggesting that regularization effects can also drive gains. Thus, biological plausibility is a sufficient but not necessary condition for effective augmentation. Practitioners should select variation levels based on their primary goal: label-preserving expansion (2–30% with ProtAug Esm) or regularization (55–77% with ProtAug GPT).

Based on these findings, we offer the following task-adaptive recommendations:

- **For resource-constrained scenarios (e.g**., ≤ **20% training data):** Use ProtAug Esm with 10% variation. This provides the most robust gains relative to both No aug and Duplicate.
- **For function- or label-sensitive tasks (most classification/regression tasks):** Use 2–30% variation with ProtAug Esm. This balances semantic preservation and diversity while maintaining label consistency.
- **For structure-related tasks or when overfitting is severe:** Consider higher variation levels (55–77%) with ProtAug GPT, which may induce regularization benefits even when label consistency is reduced.
- **When computational efficiency is a primary concern:** Random or PAM-based substitution may suffice, as they sometimes approach ProtAug performance while being substantially faster.

These findings advocate for ProtAug as a practical and cost-effective paradigm for enhancing protein sequence modeling in task-specific or resource-constrained scenarios. More broadly, our investigation highlights that the relationship between semantic plausibility and augmentation effectiveness is more nuanced than previously assumed, with both label-preserving and regularization-based mechanisms contributing to downstream improvements.

## Appendix A Statistics of Datasets

Statistics of datasets are provided in Table 3.

**Table 3.** Statistics of the datasets.

| Dataset | Function_MF | Stability | Epitope | Family | Superfamily | Fold | Secondary Structure |
| --- | --- | --- | --- | --- | --- | --- | --- |
| #. Training | 32421 | 25112 | 533 | 12312 | 12312 | 12312 | 8678 |
| #. Validation | 3587 | 6278 | 153 | 736 | 736 | 736 | 2170 |
| #. Testing | 1137 | 2413 | 78 | 1272 | 1254 | 718 | 513 |
| #. Labels | 677 | 1 | 2 | 3888 | 1961 | 1195 | 3 |
| Avg. length | 554 | 446 | 572 | 167 | 168 | 160 | 256 |

## Appendix B Fine-tuning Details

### Function Prediction

Following the TEMPROT workflow, we first generate sequence slices with a sliding window of size 500 amino acids (aa.). Additional slices are created if two consecutive slices have at least 250 aa. overlapping. Each slice retains the same function labels as the original protein sequence. Then we pass each slice through the pre-trained pLM to extract embeddings. Although in the main results we only report the performance of only training the prediction heads by freezing the pLM backbones for all tasks, we also perform full-parameter fine-tuning on all protein slices with their labels. Full-parameter fine-tuning results of three tasks are provided in Session F. Then the embeddings are derived from the [CLS] token of the last encoder block of the transformer architecture. For each protein, the embeddings from all its slices are aggregated into a single representation by taking the mean of the embeddings from all slices belonging to the same protein. After generating the combined embeddings for each protein, the next step is to train a two-layer MLP metaclassifier with binary cross entropy loss to make the final predictions. The architecture consists of one hidden layer with 1000 neurons and a ReLU activation function. The Adam optimizer is chosen and initial learning rate is set to 1e-3. The model is trained for up to 1000 epochs with a batch size of 32. The training incorporates early stopping with a patience of 15 epochs and reduces the learning rate on a plateau with a patience of 5 epochs.

### Stability Prediction

The slicing strategies for embedding generation and the meta-classification procedure for final prediction are also applied on predicting protein thermostability. The meta-classifier uses a three-layer MLP regression head with 500 and 50 neurons in the hidden layers and uses mean square error loss. The Adam optimizer is chosen, the initial learning rate is set as 1e3, and a batch size of 32 is chosen. Other training strategies are the same as those in function prediction.

### Epitope Prediction

For each peptide record, we create a segment with a maximum length of 500 aa. which is centered around the peptide sequence as much as possible. For peptides that are near the boundaries of the protein, we still make sure a segment length of 500 aa. although in this case the peptide is not at the center of the segment. If the protein sequence is shorter than 500, the entire protein sequence is used as the segment. For each segment, the original epitope binary label is retained as the label. Then we calculate the start and end positions of these peptides in the corresponding newly created protein segments. When preparing inputs for fine-tuning, we manually modify the token type ids to indicate the positions of peptides in the input segments. Specifically, in the token type ids vector only positions that belong to the peptide are set to 1 and the rest are set to 0. We extract only the residue embeddings after the pLM forward process and then take the mean of these embeddings as a single representation for a peptide in the context of its segment. No meta-classification training is used. We define a three-layer MLP classification head with 256 and 32 neurons in the hidden layers and use binary cross entropy loss for end-to-end training. The Adam optimizer is chosen, the learning rate is set as 1e-4, and a batch size of 4 is chosen. Other training strategies are the same as those in function prediction.

### Remote Homology Detection on Family, Superfamily, and Fold Levels

The slicing strategies for embedding generation and the meta-classification procedure for final prediction are also applied on Remote homology detection. The meta-classifier uses a two-layer MLP classification head with 1200 neurons and uses categorical cross entropy loss. The Adam optimizer is chosen, the learning rate is set as 1e-4, and a batch size of 32 is chosen. Other training strategies are the same as those in function prediction.

### Secondary Structure Prediction

Because each amino acid in the protein sequence has a categorical label, which indicating one of the three possible local conformation of the current position, we adopt a three-layer MLP classification head with 512 and 256 neurons in the hidden layers to predict the secondary structure label for each position in the protein sequence and use cross entropy loss. The Adam optimizer is chosen, the learning rate is set as 1e-3, and a batch size of 16 is chosen.

## APPENDIX C Details of Evaluation Metrics Definition of Fmax Score in Function Prediction

The Fmax score is defined as follows:

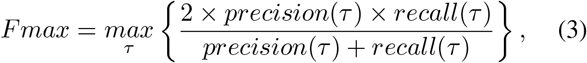

where *τ* is a threshold to be determined to maximize the F1 score. The precision and recall for a multi-label task are defined as:

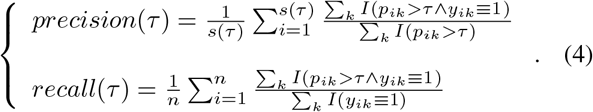

Here *s*(*τ* ) denotes the number of proteins that are predicted with at least one function, *k* is the total number of labels, *p*_*ik*_ is the predicted score for the function, and *y*_*ik*_ is the ground truth with 1 indicating the existence of the function and 0 otherwise. *n* is the total number of proteins in the evaluation.

### Definition of Repetition Ratio of Sequences

The repetition ratio of one sequence is defined as follows:

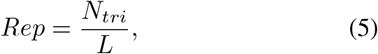

where *N*_*tri*_ is the number of triplets of identical amino acids and *L* is the length of the amino acid sequence.

The overall repetition ratio of *N* sequences is the average of the repetition ratios of these sequences:

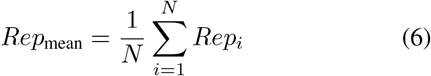

### Definition of Amino Acid Diversity

We use Shannon diversity to measure amino acid diversity in a sequence based on the occurrences of amino acids, defined as:

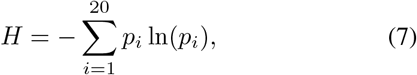

where (*p*_*i*_) is the number of occurrences of the *i* amino acid.

## Appendix D Results of Full-parameter Fine-tuning

The results of the full-parameter fine-tuning with augmented data are shown in Figure 8. Results are less table compared to fix-backbone fine-tuning, suggesting potential overfitting introduced by full-parameter fine-tuning.

**Figure 8.**
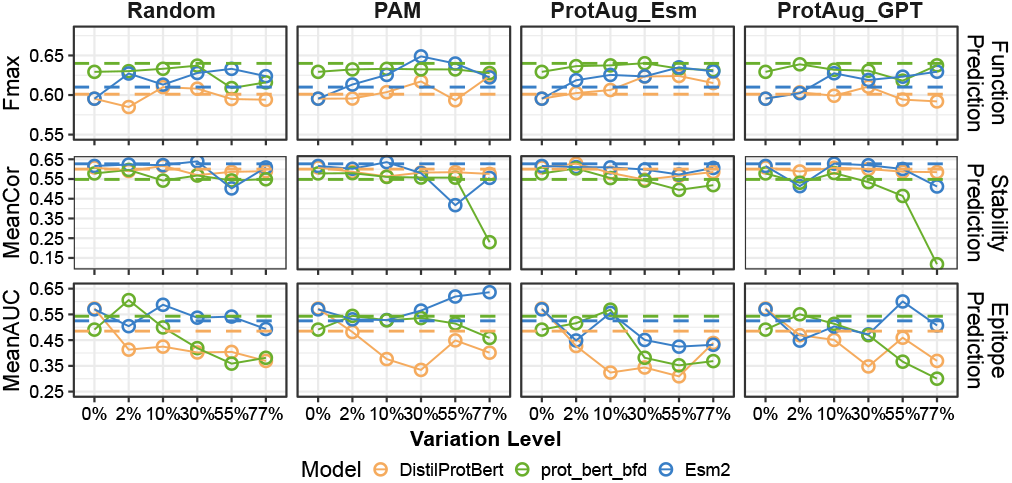
Performance of full-parameter fine-tuning pLMs with original and augmented data combined.

## APPENDIX E Ranking of Methods across various Low-Resource Scenarios

Figure 9 shows detailed rankings of different methods finetuned on 1%-10% and 20%-50% of the training data.

**Figure 9.**
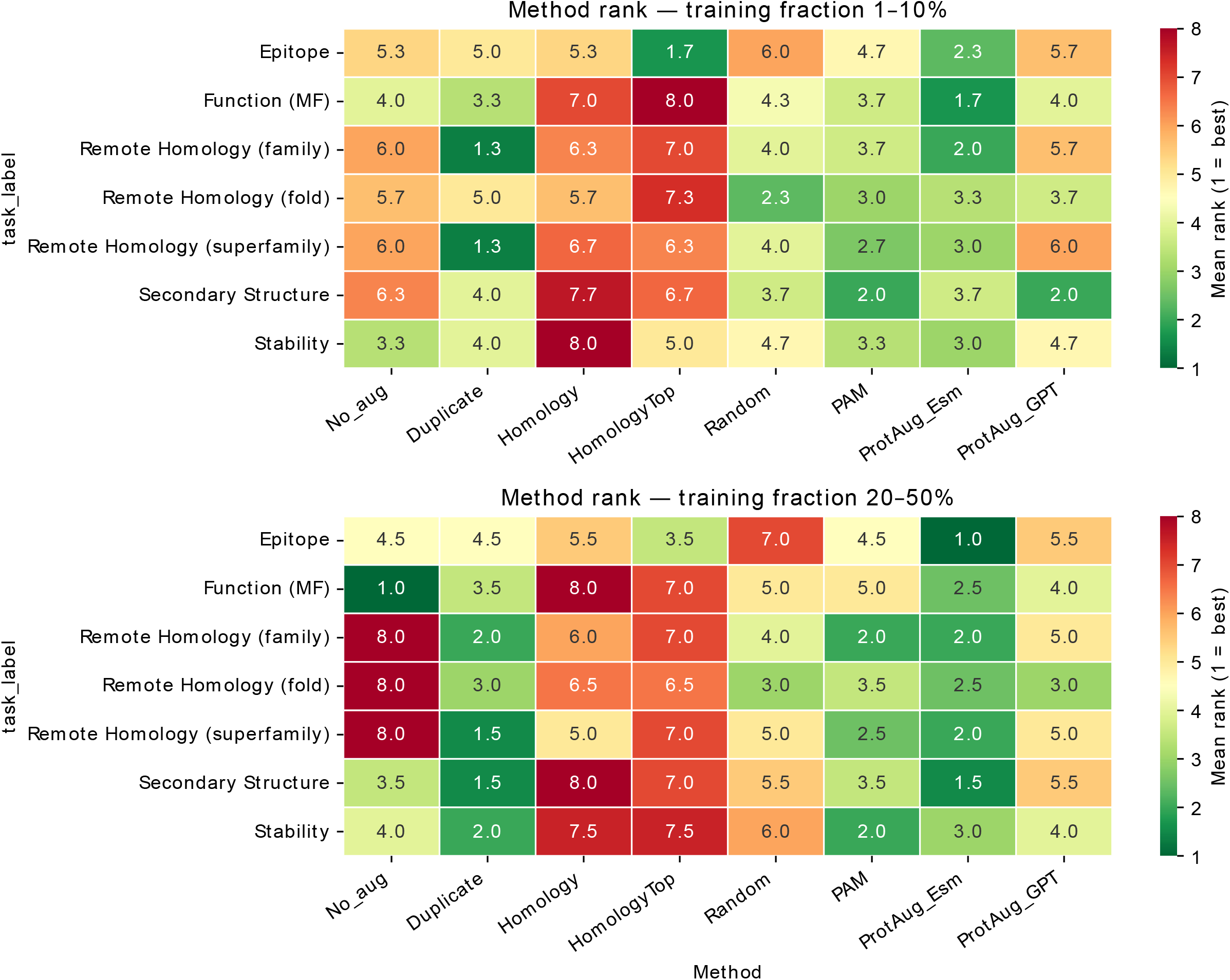
rankings of different methods finetuned on 1%-10% and 20%-50% of the training data.

## Appendix F Overall Performance on Variation Level

We perform all augmentation strategies on three pLMs across seven tasks. Results are shown in Figure 10. Generally, substitution at 2-30% variation levels yields the best results, with Esm2 consistently performs better except in Epitope prediction, where Esm2 performs the worst.

**Figure 10.**
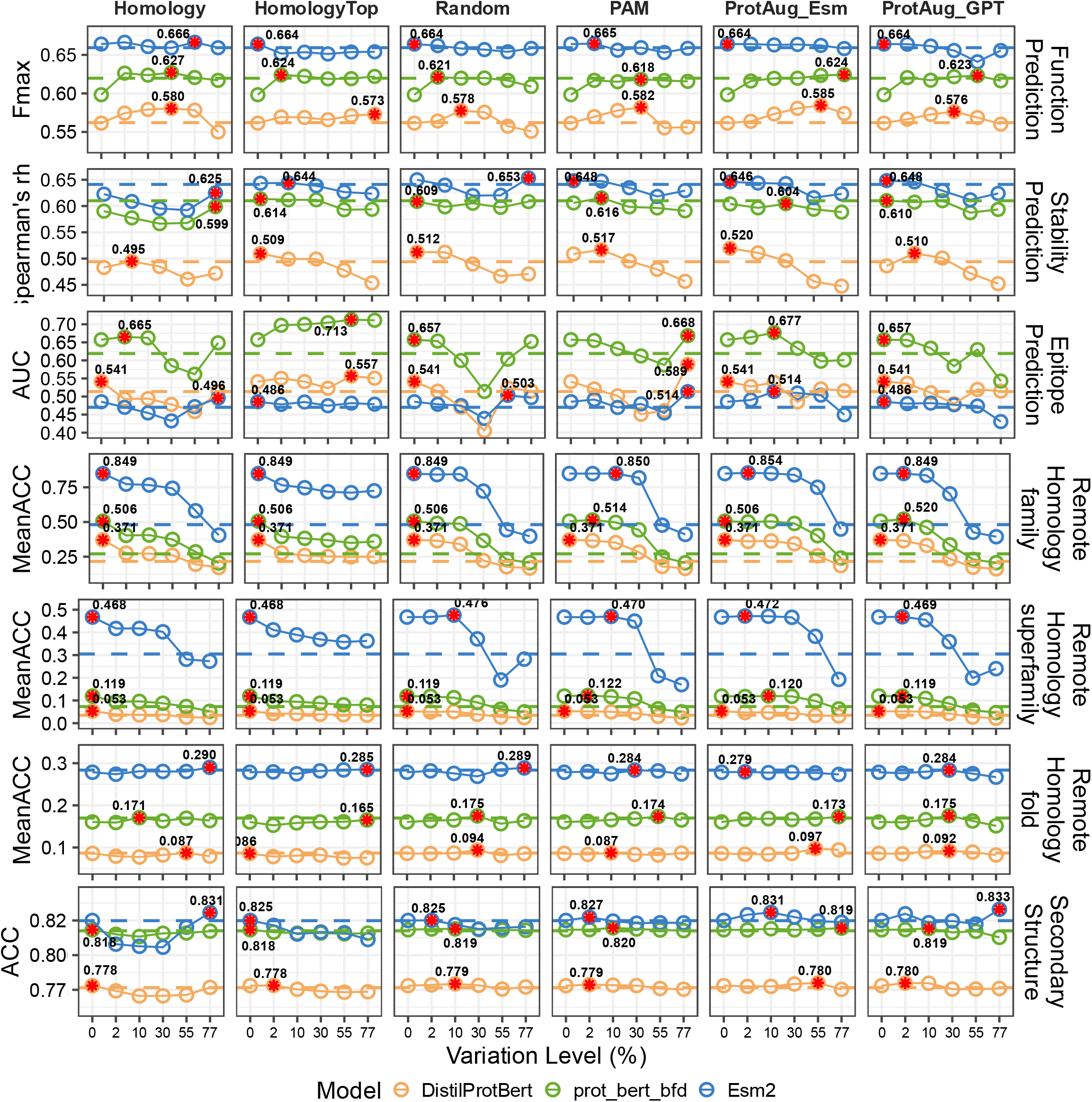
Performance of fine-tuning pLMs across variation levels. Dashed lines indicate the performance when using no augmentation. Stars denote the highest scores of corresponding models.

